# Evaluating the Evaluation of Cancer Driver Genes

**DOI:** 10.1101/060426

**Authors:** Collin J. Tokheim, Nickolas Papadopoulis, Kenneth W. Kinzler, Bert Vogelstein, Rachel Karchin

**Affiliations:** Department of Biomedical Engineering, Institute for Computational Medicine, Johns Hopkins University, Baltimore, MD 21218, USA; Ludwig Center and the Howard Hughes Medical Institute, Johns Hopkins Medical Institutions, Baltimore, MD 21231, USA; Department of Oncology, Cancer Biology Program, Johns Hopkins Medical Institutions, Baltimore, MD 21231, USA

**Keywords:** cancer genomics, DNA sequencing, driver genes, cancer mutations, computational method evaluation

## Abstract

Sequencing has identified millions of somatic mutations in human cancers, but distinguishing cancer driver genes remains a major challenge. Numerous methods have been developed to identify driver genes, but evaluation of the performance of these methods is hindered by the lack of a gold standard, i.e., bona fide driver gene mutations. Here, we establish an evaluation framework that can be applied when a gold standard is not available. We used this framework to compare the performance of eight driver gene prediction methods. One of these methods, newly described here, incorporated a machine learning-based ratiometric approach. We show that the driver genes predicted by each of these eight methods vary widely. Moreover, the p-values reported by several of the methods were inconsistent with the uniform values expected, thus calling into question the assumptions that were used to generate them. Finally, we evaluated the potential effects of unexplained variability in mutation rates on false positive driver gene predictions. Our analysis points to the strengths and weaknesses of each of the currently available methods and offers guidance for improving them in the future.

**Significance:** Modern large-scale sequencing of human cancers seeks to comprehensively discover mutated genes that confer a selective advantage to cancer cells. Key to this effort has been development of computational algorithms to find genes that drive cancer, based on their patterns of mutation in large patient cohorts. However, since there is no generally accepted gold standard of driver genes, it has been difficult to quantitatively compare these methods. We present a new machine learning method for driver gene prediction and a rigorous protocol to evaluate and compare prediction methods. Our results suggest that most current methods do not adequately account for heterogeneity in the number of mutations expected by chance and consequently have many false positive calls. The problem is most acute for cancers with high mutation rates and comprehensive discovery of drivers in these cancers may be more difficult than currently anticipated.

## Introduction

The search for genetic drivers of cancer has rapidly progressed with systematic exome sequencing studies ^1^. A major goal of these studies is to identify signals of positive selection and distinguish them from random accumulation of passenger mutations. The first exomic analyses attempted to identify candidate driver genes as those having more mutations than expected from some presumed background somatic mutation rate, corrected for base context, gene size, and other variables ^2–4^. Subsequent work has considerably refined the variables involved in determining whether a gene is more mutated in cancers than expected by chance. This has led to a variety of “significantly mutated gene” (SMG) methods that adjust for covariates such as replication timing and gene expression as well as including more sophisticated metrics of mutational base contexts^5,6^.

An alternative approach to finding cancer drivers employs ratiometric methods. Rather than attempting to determine whether the observed mutation rate of a gene in cancers is higher than expected by chance, these methods simply assess the composition of mutations normalized by the total mutations in a gene. The ratiometric 20/20 rule ^7^ evaluates the proportion of inactivating mutation and recurrent missense mutations in a gene of interest. Other ratiometric approaches use mutation functional impact bias ^8^, mutational clustering patterns ^9,10^ and mutation composition patterns Here, we describe a new machine learning-based, ratiometric method (20/20+) that formalizes and extends the original 20/20 rule and enables automated integration of multiple features of positive selection.

Rigorous and unbiased evaluation is necessary to inform users about the comparative utility of prediction methods, including the new method described herein. In many investigative domains, there is a generally accepted gold standard against which predictions can be benchmarked. However, only a limited number of genes have been fully vetted as cancer drivers. In previous work, driver prediction has been benchmarked by significant overlap with the Cancer Gene Census (CGC) ^11^, which is a manually curated list of likely but not necessarily validated drivers ^8,9,12^, by agreement with a consensus gene list of drivers predicted by multiple methods ^13^, and by number of “suspicious” predicted driver genes with no clear relevance to cancer ^5^. To our knowledge, a systematic framework for the evaluation of somatic mutations that can be generally applied has not been previously developed.

In this work, we present a framework for such evaluations. The framework has five components, some of which have been previously applied in isolation, but not as part of a unified system. We considered overlap with Cancer Gene Census (CGC), agreement between methods, comparison of observed vs. theoretical p-values, number of significant genes predicted, and prediction consistency on independent partitions of each data set. To implement this framework, we first collected 729,205 published somatic mutations from 34 cancer types ^10,14^ (Figure 1). These mutations were composed of single base substitutions and in-frame and out-of-frame insertions and deletions (indels) of less than 10 bp. We then compared various methods on the full pan-cancer set and on four selected cancer types with diverse mutation rates-pancreatic adenocarcinoma (PDAC), breast adenocarcinoma (BRAC), head and neck squamous cell carcinoma (HNSCC), and lung adenocarcinoma (LUAD).

**Figure 1.**
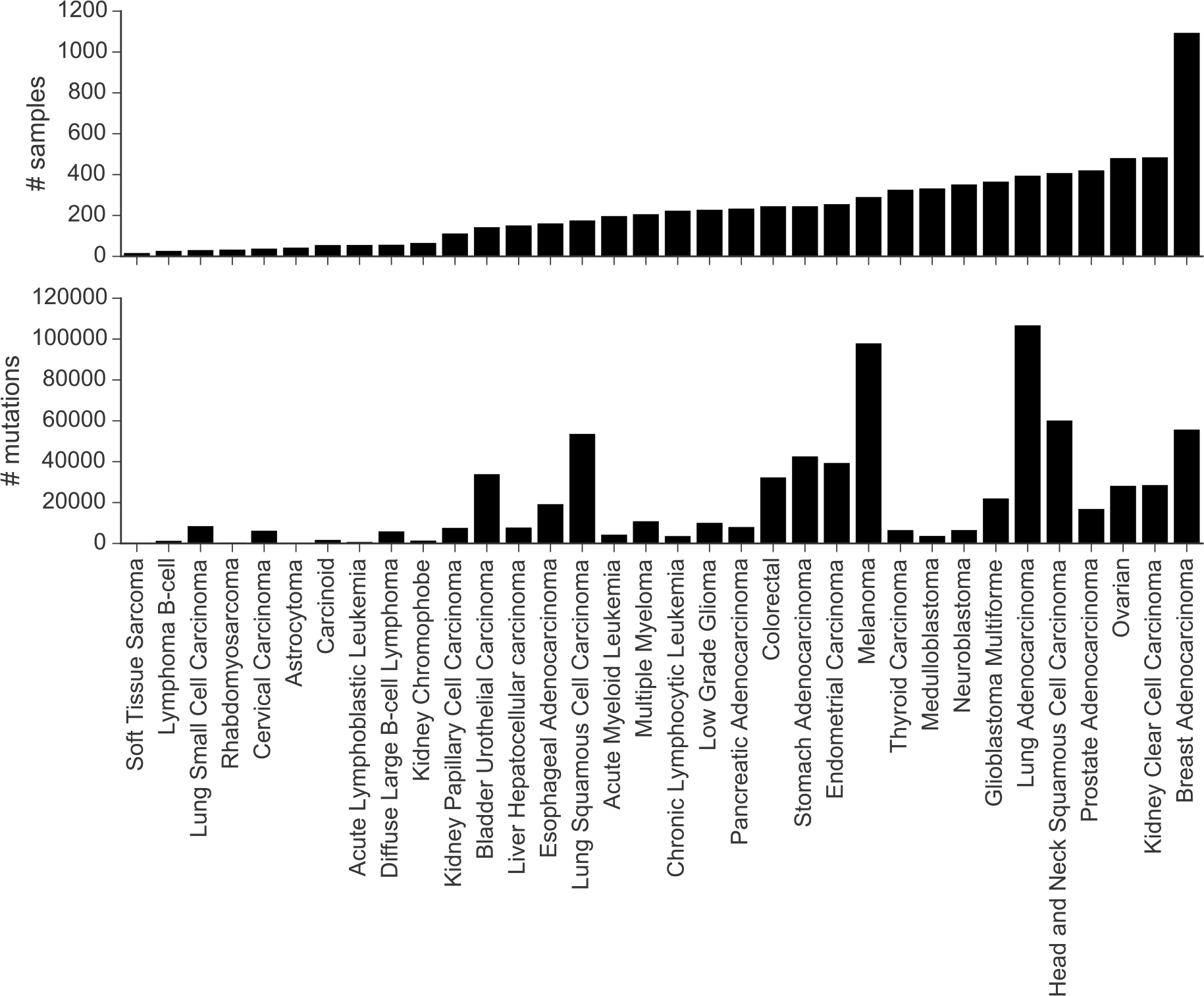
Summary of evaluation dataset. The evaluation dataset consisted of mutations spanning 34 cancer types. All included mutations were small somatic variants. Cancer types are ordered from left to right by number of samples, ranging from for soft tissue sarcoma to 1,093 for breast adenocarcinoma, with an average of 232 samples per cancer type. These cancer types span a wide range of solid and several liquid cancers, including multiple tissues and cell-types or origin, different background mutation rates and different numbers of available samples. For each cancer type, total mutations and number of available samples are shown.

## Results

### Overlap of the driver genes predicted by each method

Eight methods were evaluated: MutsigCV ^14^, ActiveDriver ^15^, MuSiC ^6^, OncodriveClust ^9^, OncodriveFM ^8^, OncodriveFML (https://bitbucket.org/bbglab/oncodrivefml), TUSON ^10^ and 20/20+ (https://github.com/KarchinLab/2020plus). All data, on all cancers, was considered (“pan-cancer”) in these comparisons. First we assessed overlap of the predicted driver genes with the CGC. We considered only those CGC genes typed as somatic, missense, frameshift, nonsense or splice site, excluding translocations, large amplifications/deletions and other mutation consequence types not addressed in our study, yielding a total of 188 CGC genes. Though the driver genes predicted by all methods were enriched for CGC genes, the predicted drivers by any individual method did not contain a majority of CGC genes (Table S2, Figure 2A).

**Figure 2.**
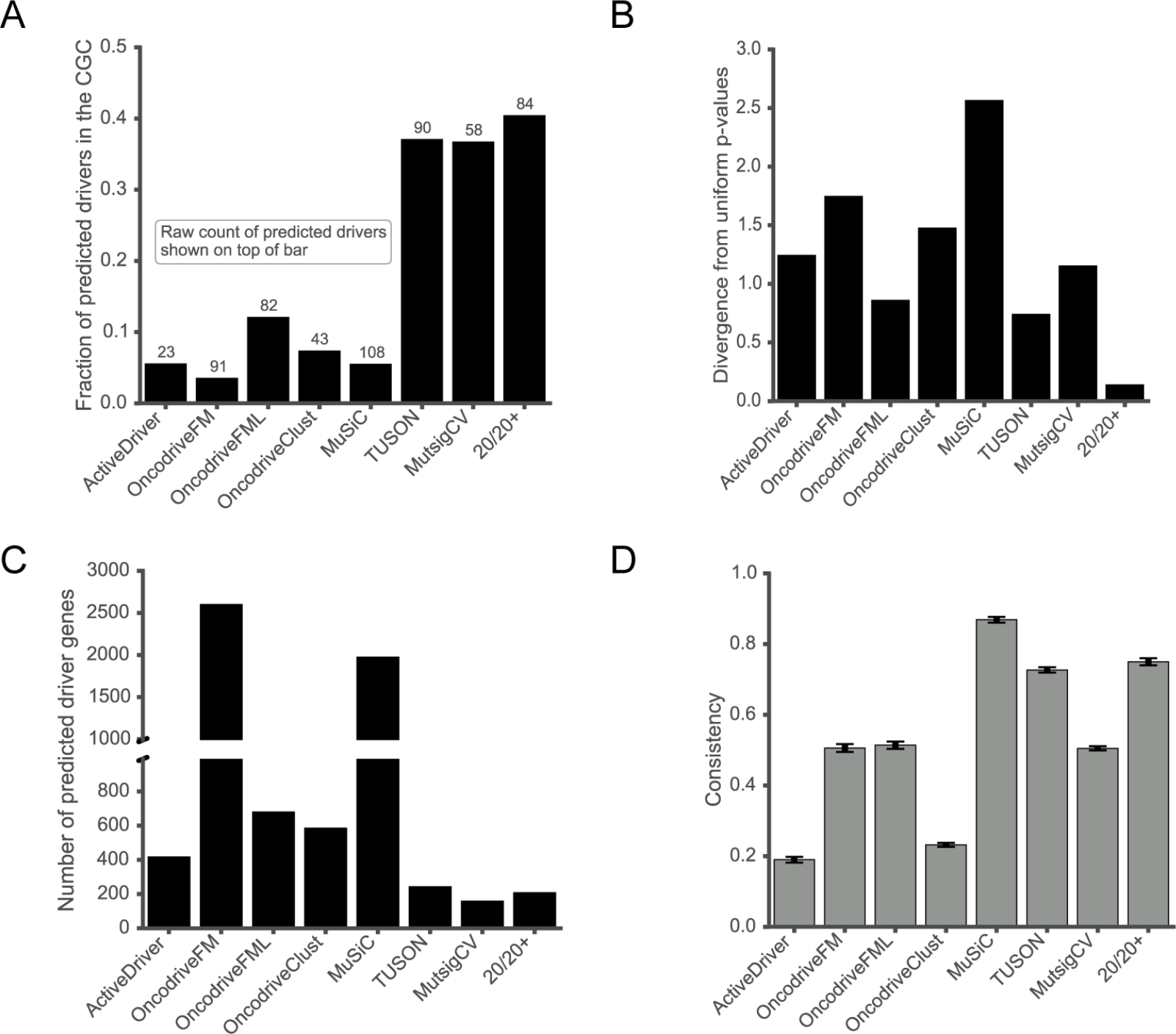
Performance of eight driver prediction methods on the five component evaluation protocol. **A.** Fraction of predicted driver genes (q≤0.1) that are found in the Cancer Gene Census (CGC) (downloaded 04/01/16). Raw count of predicted driver genes indicated on top of each bar. **B.** Divergence from uniform p-values, measured as mean log fold change (MLFC) between a method’s observed and desired theoretical p-values. **C.** Number of predicted driver genes. Driver gene is defined as having Benjamini-Hochberg adjusted p-value q≤0.1. **D.** Consistency of each method measured by TopDrop Consistency (TDC) at depth of 100 in the method’s ranked list of genes. Error bars indicate ±1 SEM (standard error of the mean) across 10 repeated splits of the data.

Genes predicted by more than one method may be more likely to be drivers ^13^. For each method, we calculated the fraction of predicted drivers that were unique, those predicted by at least one, two, or three other methods (Figure S1, Dataset S1). As shown in Figure S1, there was little consensus in prediction of driver genes among the methods. The majority (59% to 80%) of genes identified by MuSiC, ActiveDriver, OncodriveClust, OncodriveFML, or OncodriveFM were *not* observed by any of the other seven methods. The fractions of genes identified by TUSON, 20/20+, and MutsigCV that were *not* identified as driver genes in at least one of the other seven methods was 14%, 19%, and 33%, respectively.

### Observed vs. expected p-values

Given the lack of agreement among these various methods, we compared p-values reported by each method to those expected theoretically. Such comparisons are often used in statistics and can indicate invalid assumptions or inappropriate heuristics. Theoretically, the p-value distribution should be approximately uniform after likely driver genes are removed ^16^. Therefore, we removed all genes predicted to be drivers by at least 3 methods after Benjamini-Hochberg multiple testing correction (q≤0.1) and any remaining genes in the CGC. We assumed that the number of bona fide driver genes not removed by this procedure would be small. To quantify the differences between the observed p-values and those expected from a uniform distribution, we developed a measure named Mean Absolute Log2 Fold Change (MLFC) (Online Methods). MLFC values near zero represent the smallest discrepancies and the closest agreement between observed and theoretical p-values.

One method (20/20+) had an MLFC that was 5-fold lower than the seven others (Figure 2B). We also compared observed and theoretical p-values with quantile-quantile plots, which provide a detailed view of p-value behavior (Figure S2A). 20/20+ p-values had by far the best agreement with theoretical expectation across the entire range of supported values. In the critical range typically used to assess statistical significance (p≤0.05), OncodriveClust, OncodriveFM, OncodriveFML, ActiveDriver and MuSiC substantially underestimated p-values, while MutsigCV substantially overestimated them (Figure S2B).

### Number of predicted driver genes

The number of predicted driver genes (q ≤ 0.1) ranged from 158 (MutsigCV) to 2600 (OncodriveFM) (Figure 2C). There were two obvious categories of methods with respect to predicted driver genes: MutSigCV, 20/20+, and TUSON predicted 158 to 243 genes, while the remaining five predicted over 400 driver genes.

### Driver gene prediction consistency

Statistical methods suffer from both systematic and random prediction errors ^17^. When no gold standard is available, it is difficult to estimate systematic error, but possible to estimate random error by measuring the variability of predictions. We tested the 8 methods on ten repetitions of a random two-way split of the all samples in our dataset, while maintaining the proportion of samples in each cancer type. An ideal method would produce the same list of driver genes, ranked by p-value, for each half of the split. For a fair comparison, we considered that methods predicting many drivers would be less likely to have consistent rankings than those predicting only a few. Thus, we developed a measure named TopDrop Consistency (TDC) (Online Methods) that examines the overlap between genes ranked at a defined depth (e.g., the top 100 genes) for each half of the random split. Examining TDC at a depth of 100 genes showed MuSiC, 20/20+, and TUSON to be the three with the highest consistency (Figure 2D). Most methods decreased in consistency when the gene depth was varied between 20 and 300, but the ordering of the TDC scores among the eight methods remained relatively stable (Figure S3).

### Evaluation of specific cancer types

To evaluate whether methods performed differently on specific cancer types, rather than on the pan-cancer dataset, we repeated our evaluations using four specific cancer types. We considered two moderate mutation rate cancers (pancreatic adenocarcinomas, with median of 0.7 mutations/MB, breast adenocarcinomas, with median of 1 mutation/MB) and two high mutation rate cancers (head and neck squamous carcinomas, with median of 3.2 mutations/MB, and lung adenocarcinomas, with median of 6.7 mutations/MB). These mutation rates were based on the same dataset used throughout this study and described above. As with the pan-cancer types, the 20/20+ method had the least discrepancy between observed and theoretical p-values (Figure S4A). For the TopDrop Consistency (TDC) evaluation, we used a rank depth of 10 genes rather than the 100 used for pan-cancer, as it is likely that this number is closer to the number of driver genes in a single cancer type. The most consistent methods were 20/20+ and MuSiC (Figure S4B) and the least consistent methods were ActiveDriver and OncodriveClust. Driver genes in head and neck squamous carcinoma (HNSCC) were the most consistently predicted overall.

The number of cancer-specific predicted driver genes varied widely (Figure S4C). PDAC had the fewest predicted drivers (q≤0.1), ranging from 3 (TUSON) to 49 (OncodriveFM), while LUAD had the most, ranging from 9 (TUSON) to 922 (OncodriveFM). ActiveDriver, OncodriveFM and OncodriveClust predicted hundreds of cancer-type specific drivers, while OncodriveFML, MuSiC, 20/20+ and MutsigCV tended predicted no more than 40 in any of the four cancer types.

### Variability in background mutation number

Because only a small fraction of the total somatic mutations in any common solid tumor affects driver genes, the remaining mutations can be considered passengers, reflecting the mutation “background”, i.e., all mutations that occurred during the divisions of the tumor cells, from embryogenesis until the tumor was surgically removed ^18^. The total number of mutations (drivers plus passengers) is therefore only slightly larger than the number of passenger mutations, and for simplicity, we refer to this number as the background mutation rate. The median background mutation rate for cancer types in our pan-cancer set varied over two orders of magnitude (Figure S5), with individual samples varying over an even larger range. Mutation rates vary among individual genes and are influenced by nucleotide context, environmental factors, gene expression, chromatin state, replication timing, DNA repair activity, strand, and perhaps by a variety of factors that have yet to be discovered ^5, 19–21^.

We analyzed the possible impact of unexplained variability in background mutation rate on expected false positive driver gene predictions. First, we applied a binomial model previously used for driver gene detection power analysis ^14^. The model assumes a gene-specific background mutation rate μ, which is set to a relatively high value, corresponding to genes in the 90th percentile of mutation rate. We used the binomial to set a critical value for driver gene prediction, i.e., the number of mutations required for a gene to be considered significantly different from the background. Next, we modeled the situation where the genes actually had mutation rates that varied around μ, using a beta-binomial model. We estimated the false positives expected under the binomial, after a highly conservative multiple testing correction (Bonferroni). The number of mutations required to meet or exceed the binomial critical value was compared with that of the beta-binomial, for different background mutation rates, levels of variability (beta-binomial coefficients of variation) and for sample size ranging up to 8,000 (Figure 3A). Levels of variability defined by coefficients of variation (CV=0.05, 0.1 and 0.2) were chosen to approximate low, medium and high unexplained variation around the mean. As the number of samples increased, so did the number of expected false positives. At the low end of background mutation rates (0.5 mutations/MB) the expected false positives remained low, even when 8,000 samples were evaluated, regardless of the level of variability. At an intermediate background mutation rate of 3.0 mutations/MB and with high unexplained variability, ~1000 false positives were expected from 8000 samples. At a high background mutation rate (10.0 mutations/MB), both medium and high unexplained variability produced many thousand expected false positives.

**Figure 3.**
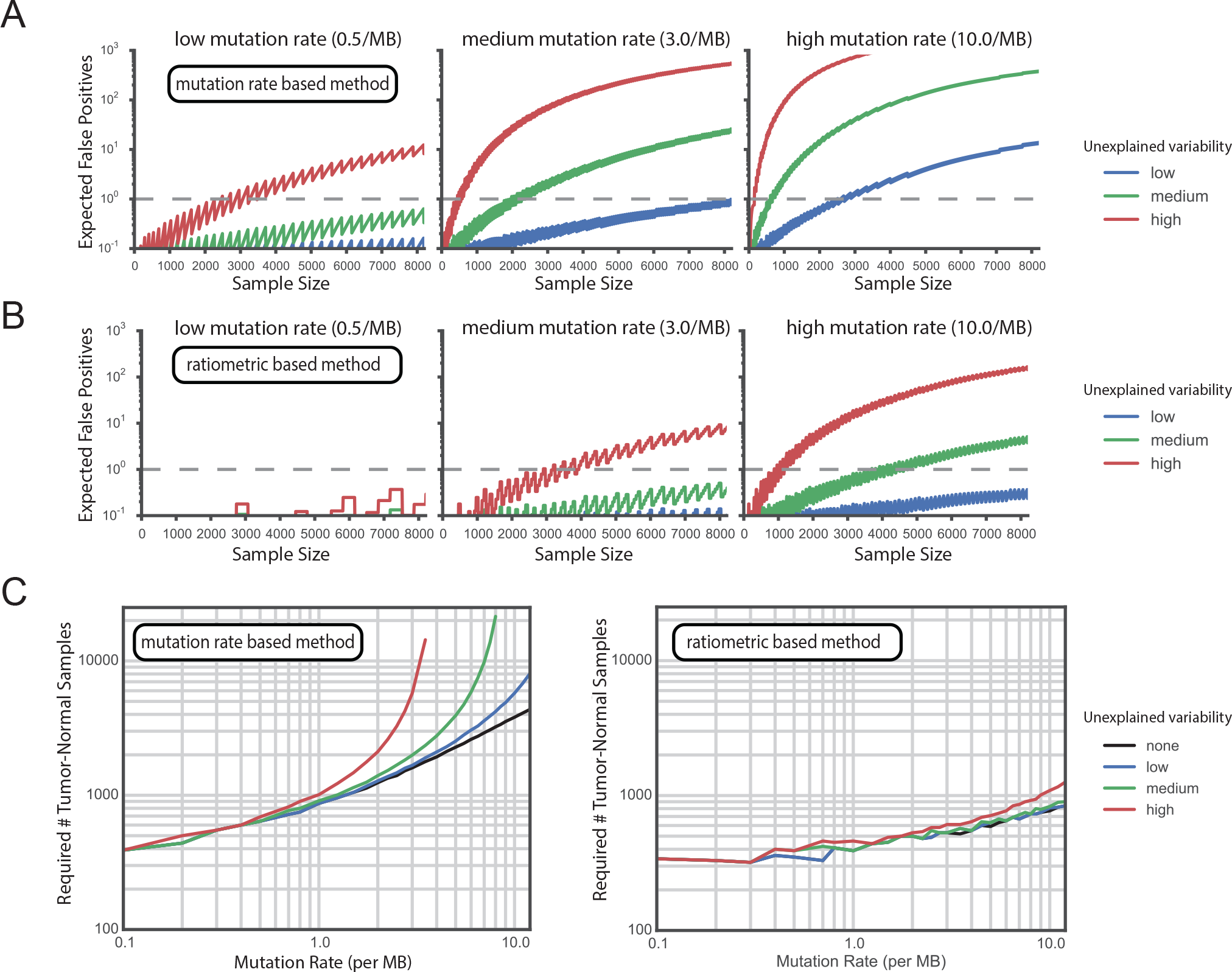
Models of mutation-rate based and ratiometric based methods suggest decrease in false positives and increased power with ratiometric approach. **A.** Expected false positives for a mutation-rate-based predictor that identifies genes with increased mutation rate over background. **B.** Expected false positives for a ratiometric predictor that identifies genes with increased inactivating mutation fraction over background. For both A and B, we assume there is unexplained variability in either background mutation rate or inactivating mutation fraction that is not accounted for in driver gene prediction. False positives are shown as a function of sample size (up to 8000 paired tumor-normal samples) for low (0.5 mutations/MB), medium (3.0 mutations/MB) and high (10.0 mutations/MB) background mutation rates and low (blue), medium (green) and high (red) unexplained variability (coefficients of variation 0.05, 0.1, and 0.2, respectively). The dashed line indicates one expected false positive. For the mutation-rate based method, the number of false positives increases to undesirable levels for high mutation rates, particularly when there is high unexplained variability. **C.** Sample size required for near-comprehensive detection of intermediate-effect driver genes (90% detection and 2% effect size/increase with respect to background). Results are shown for scenarios with no unexplained variability (black), low (blue), medium (green) and high (red) unexplained variability (coefficients of variation 0.0, 0.05, 0.1, and 0.2, respectively). The number of required samples for the mutation-rate based method becomes very large for moderate to high mutation rates and levels of unexplained variability, but it is considerably lower for the ratiometric method. MB=mega base.

We reasoned that unexplained variability might also have an impact on power calculations to estimate how many samples must be sequenced to find the majority of cancer driver genes. To this end, we repeated previous calculations performed with a binomial power model, in which the required sample size was estimated to be 600-5,000 per cancer type ^14^. The original model was parameterized to detect intermediate frequency driver genes, having 2-20% mutation rates above background per sample, with background defined by genes in the 90th percentile of background mutation rates. First, we calculated the sample size required to detect 90% of these drivers, given exome-wide backgrounds of 0.1-10 mutations / MB, and a conservative estimate of 2% effect size (Online Methods). Next, we calculated the sample size required if the gene mutation rate varied around the original estimate, using a beta-binomial model with different coefficients of variation (CV=0.05, 0.1, 0.2). The binomial power model was in accord with previous estimates. However, when unexplained variability was taken into account, the number of required samples increased sharply, particularly for higher background mutation rates (Figure 3C).

### Variability in ratiometric features

Ratiometric features would appear to have significantly less variability than background mutation rates. They are typically defined as either a fraction of a specific mutation type out of the total mutations in a gene, or a ratio of one mutation type to another. Figure S5 shows the variability of the median ratio of non-silent to silent mutations for cancer types in our pan-cancer set. Rather than varying over orders of magnitude, this feature varies over one or two units and individual samples vary over less than ten units. However, ratiometric features might also be sensitive to unexplained variability in their background distributions. Therefore, we calculated the expected false positives and statistical power of a simple ratiometric feature, the fraction of mutations in a gene having a specific non-silent mutation consequence type. A slightly modified version of the calculations used for background mutation rate was applied (Materials and Methods). We observed improved false positive control and reduction in the number of required samples, particularly for high mutation rate cancers (Figure 3B, 3C).

## Discussion

A major goal of the huge public investment in large-scale cancer sequencing has been to find the majority of driver genes. Robust computational prediction of drivers from small somatic variants is critical to this mission, and it is essential that the best methods be identified. While many such methods have been proposed, it has been difficult to evaluate them because there is no gold standard to use as a benchmark. Here we developed an evaluation framework for driver gene prediction methods that does not require a gold standard. The framework includes a large set of small somatic mutations from a wide range of cancer types and five evaluation metrics. We propose it can be used to systematically evaluate new prediction methods and compare them to existing methods. The results would be more informative to users of these methods than current ad hoc approaches.

To apply the framework to a new method, a ranked list of predicted driver genes can be generated from the pan-cancer mutation dataset (Figure 2), including a p-value and a Benjamini-Hochberg corrected q-value for each gene. The choice of a threshold q≤0.1 to define driver genes worked well in our evaluations, but can be adjusted if so desired. The same threshold should be used for fair comparison of different methods.

By calculating the overlap fraction of predicted drivers with both the Cancer Gene Census (CGC) and the eight methods evaluated here, it is possible to quickly determine whether a new method is on the right track. Two baselines of good performance are a method’s ability to recapitulate many of the well-studied cancer genes in CGC and ability to identify a core set of genes that are predicted as drivers by several other methods. Here the methods with strongest support by these criteria were 20/20+, TUSON and MutsigCV. Roughly 40% of predicted drivers by these methods were in CGC, contrasted with roughly 10% of predicted drivers by the remaining methods. They also had substantially more overlap with other methods and predicted the smallest fraction of unique genes, those having no overlap with other methods (Figure 2A, 2B). While detecting some unique genes is desirable for purposes of discovering novel drivers, if the fraction of unique predictions is much greater than half, a method may be prone to false positives. Comparing the gene p-value distribution of a method of interest with theoretically expected p-values and with the total number of predicted driver genes may help make sense of such a result.

The TopDrop Consistency metric (TDC 100 and TDC 10) evaluates the stability of the top *k* genes in a ranked list, when a method is applied repeatedly to matched random partitions of a dataset. The most consistent methods evaluated here had a high percentage of the top 100 genes consistently ranked on partitions of the pancancer dataset: MuSiC (87%), 20/20+ (75%) and TUSON (73%). These three methods also consistently ranked >80% of the top 10 genes in cancer-specific data sets of breast adenocarcinoma (BRAC), lung adenocarcinoma (LUAD) and head and neck squamous cell carcinoma (HNSCC) (except for TUSON on LUAD) (Figure S4B). We suggest that the ability to reproduce approximately three-quarters of the same top-ranked genes is a reasonable baseline standard.

Based on the assumption that the total number of cancer driver genes is very small when compared to the total number of human genes ^7^, it is reasonable to assume that p-values assigned to all genes should be approximately uniform after a core set of well-established driver genes have been dropped. The Mean Log Fold Change (MLFC) metric proposed here quantifies the deviation between these theoretically expected p-values and the observed p-values generated by a method. Lower values of MLFC are desired; 20/20+ had the lowest MLFC of methods evaluated, both for pan-cancer and for all four cancer types evaluated (Figure 2B and Figure S4C). The highest MLFC values were for MuSiC, OncodriveClust and ActiveDriver. More detailed p-value behavior can be seen in quantile-quantile plots that compare the observed p-values with a theoretical uniform distribution (Figure S2A, S2B). The plots show that MuSiC, ActiveDriver, OncodriveClust, OncodriveFM and OncodriveFML p-values are underestimated (lower than theoretically expected) in the critical range (p-values from 0 to 0.05) while the p-values of TUSON and MutsigCV are systematically overestimated. The low-end of the p-value range is critical because it corresponds to the threshold used to call driver genes (q=0.1 in this work). If p-values are underestimated in this range, too many genes will be called as drivers. In fact, the methods that underestimate p-values predict the largest number of drivers and have the highest fraction of uniquely predicted drivers (Figure 2C, Figure S1).

These results suggest that the true number of driver genes represented in the pan-cancer dataset is closer to the number predicted by 20/20+ (208) and TUSON (243), which have the lowest MLFC. If a new method is tested and reports a substantially larger number of drivers than these two methods, the MLFC and quantile-quantile plots should be carefully checked for discrepancies with theoretically expected p-values.

The MFLC also has substantial implications for the accuracy of driver gene prediction methods. The relatively high MFLC of several methods brings into question the validity of the assumptions or analytic methods used in their construction. We believe that the most likely problem is with the assumptions rather than the analytic methods, which all appear to be well thought-out. And the most likely problem with the assumptions is that there is unexplained variability in the background mutations rates. This variability may be tumor-type specific or even patient-or tumor-specific. In an effort to understand the potential effects of such unexplained variability on cancer gene driver predictions, we modeled the situation encountered when various numbers of mutations were available for evaluation. The results were striking in that high levels of variability produced huge numbers of false positives when background mutations rates were high. Thus, cancers with high background mutation rates (such as those associated with environmental carcinogens) are the most problematic for driver prediction methods. This analysis also demonstrated how unexplained variability in gene mutation rates might confound power calculations. Estimates of near-saturation discovery of drivers using ~5,000 samples in high mutation rate cancers, such as melanoma ^14^, are likely to be overly optimistic, and such discovery may require more resources than currently projected. Identifying the optimum driver gene prediction methods will be an important part of this effort, and we believe that the evaluation protocol described here will help to test and improve those methods.

## Materials and methods

### Data collection

The pan-cancer dataset consists of 729,205 small somatic variants (SSVs) encompassing 7,916 distinct samples from 34 specific cancer types by merging data in published whole-exome or whole-genome sequencing studies utilized by TUSON (http://elledgelab.med.harvard.edu/wpcontent/uploads/2013/11/Mutation_Dataset.txt.zip)^10^ and Mutsig (http://www.tumorportal.org/load/data/per_ttype_mafs/PanCan.maf) ^14^ and removing duplicate samples in both studies. Any studies that did not report silent mutations were removed. Data in ^10,14^ originated from The Cancer Genome Atlas (TCGA), International Cancer Genome Consortium (ICGC), or the Catalogue of Somatic Mutations in Cancer (COSMIC) database ^22^. We further applied quality control to this data by filtering out hyper-mutated samples (>1,000 intra-genic small somatic variants) ^7^, and regions prone to mutation calling artifacts (any sequencing read mappability warning catalogued in the UCSC Genome Browser ^23^). The CRAVAT webserver (version 3.0) ^24^ was used to automatically retrieve the mappability warning codes. Gene names were standardized to HUGO Gene Nomenclature Committee through converting previous symbols and synonyms to the accepted gene name (downloaded 1/29/2015:ftp://ftp.ebi.ac.uk/pub/databases/genenames/locusgroups/protein-codinggene.txt.gz). Four cancer-specific mutation sets were extracted from the pan-cancer set: pancreatic adenocarcinoma (PDAC), breast adenocarcinoma (BRAC), lung adenocarcinoma (LUAD) and head and neck squamous carcinoma (HNSCC).

#### Mean Log Fold Change

The Mean absolute Log Fold Change (MLFC) is a metric of discrepancy between an observed p-value distribution reported by a method and a theoretical uniform null distribution. We define *P*(*i*) = *i’th smallest p – value* and 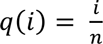, then

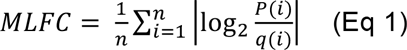

where P represents the observed p-value, q is the corresponding expected p-value from a uniform distribution, n is the total number of genes, and MLFC is the average difference of observed and theoretical p-values. Values of MLFC near zero indicate smaller discrepancies, and therefore better statistical modeling of the passenger gene null distribution. To evaluate TUSON, which reports both an oncogene and tumor suppressor gene p-value, we calculated the average MLFC score between the two.

#### TopDrop Consistency

Consistency assesses stability in gene ranking. Each method was applied to 10 repeated random splits, consisting of two disjoint halves of the full data. For pan-cancer assessment, the proportion of samples from each cancer type was maintained in each half. Disjoint halves were scored separately by each method, and genes were ranked from low to high p-values. For a fair comparison between methods, we considered a specific depth of top-ranked genes, rather than a fixed q-value threshold. This is because consistency becomes harder to achieve as the number of top-ranked genes gets larger. For example, a method that predicts 100 significant genes at q≤0.1 has an advantage in consistency over a method that predicts 1000 significant genes at that threshold.

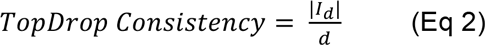

where d is the designated depth of interest for the ranked gene list and *I_d_* is the *TopDrop Intersection* (Eq 3).

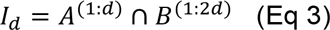

defined as the intersection between predictions from the two random halves “A” and “B” such that the top d genes in “A” do not fall past twice the designated depth (2d) in “B”, and d is the depth of interest in the ranked gene list.

We expect that all methods will lose statistical power and have greater random sampling error when they are predicting on a dataset that has been split in half. Therefore, we chose to allow genes to fall twice as far down the list in the “B” half of the split, to better distinguish random effects and methods with intrinsically low consistency.

### 20/20+: a new method for driver gene prediction

20/20+ is designed to advance and generalize the 20/20 rule ^7^, a method to identify tumor suppressor genes (TSGs) and oncogenes (OGs) using ratiometric features based on somatic mutations in the COSMIC database (cancer.sanger.ac.uk) ^25^. The core elements of the 20/20 rule are 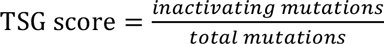 and 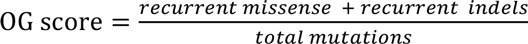. For each gene, the two scores are calculated, and a decision tree process is applied (Figure S6) by considering a series of fixed thresholds on the TSG score, OG score, the inactivating mutation count (numerator of TSG score) and the recurrence count (numerator of OG score). The thresholds were manually selected, based on extensive manual curation and tuned to the composition of COSMIC in the year when the rules were designed.

Our goal was to generalize 20/20 by replacing the decision tree and fixed thresholds with a more flexible learning framework. We selected a Random Forest (ensemble of decision trees) and used the set of oncogenes (OGs) and tumor suppressor genes (TSGs) identified by the original 20/20 rule as a training set. The Random Forest learns three classes of genes: TSGs, OGs and passengers. Driver predictions include either TSGs or OGs.

#### Predictive features

We designed a set of 24 features (Table S1). Many of the features are components of the 20/20 rule OG and TSG scores, and we included several ratiometric features not in the original 20/20 rule, *e.g.,* ratio of missense to silent mutations, as well as features that represented mutation functional impact and gene importance. Normalized missense entropy, a measure of positional clustering, was calculated as

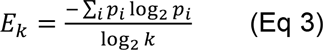

where *k* is the total number of missense mutations in a gene and *p_i_* = (count of missense mutations in the *i^th^* codon) / *k*.

We calculated inactivating single nucleotide variant (SNV) fraction p-value, normalized missense entropy p-value and missense VEST score p-value (Table S1) with Monte Carlo simulations, to approximate their null probability distributions. Briefly, for each gene, SNVs were moved with uniform probability to any matching position in the gene sequence, holding the total number of SNVs fixed. A matching position was required to have the same base context (C*pG, CpG*, TpC*, G*pA, A, C, G, T) ^26^ as the observed position. This method of generating a null distribution controls for the particular gene sequence, gene length and mutation base context. The number of SNVs remains the same, but the mutation consequence of a SNV may change. For example, a SNV that generates a missense mutation may generate a nonsense mutation in its new position.

Based on the Random Forest feature importance metric (decrease in Gini index) ^27^, the four most informative features were inactivating SNV fraction p-value, frameshift indel fraction, nonsense fraction and missense fraction. A complete list of feature rankings is shown in Figure S7.

#### Random Forest

20/20+ utilizes a three-class Random Forest ^27,28^ machine learning algorithm from the *random Forest* R package to predict whether a gene is an oncogene (OG), tumor suppressor gene (TSG), or passenger gene. With only 54 OGs and 71 TSGs labeled by the original 20/20 rule, the number of passenger genes far exceeds the number of labeled driver genes, creating a problematic class imbalance. We employ a subsampling approach ^29^, in which passenger genes are sampled at a 1:1 ratio to oncogenes plus TSGs. To compensate for the remaining oncogene and TSG imbalance, the Random Forest is trained with class weights inversely proportional to the sampled frequency of the class.

Predictions were made with a random forest of 200 trees. For the pan-cancer data, we used 10-fold gene hold-out cross-validation to avoid overfitting ^30^. The procedure of 10-fold cross-validation was repeated 5 times, and the resulting scores from each gene were averaged to limit minor fluctuations in scores due to randomization in the cross-validation folds. Each gene was scored as the fraction of trees that voted for OG, TSG, or passenger gene. A driver score for each was calculated as the sum of the OG and TSG scores.

The statistical significance of each gene score was computed with an extension of the Monte Carlo simulation algorithm described above. Since mutations that result in insertions and deletions will not change their mutation consequence type by being randomly moved to another nucleotide in the same gene, they were moved to a random position in a different gene. This gene was selected based on a multinomial model, with probability proportional to the CDS length of the originating gene. For each gene, the Monte Carlo simulation was repeated 10 times, and for each simulation all 24 features were computed. In this process, protein interaction network features (degree and betweenness) were, additionally, permuted as a pair. The features of gene length, replication timing, HiC value, and average Cancer Cell Line Encyclopedia (CCLE) gene expression were not altered. Next, each “simulated” gene sequence was scored with the Random Forest previously trained on the real data. The resulting OG, TSG and driver scores for all genes were utilized as an empirical null distribution. To compute a p-value for a gene score, we used the fraction of simulated genes with a score equal to or greater than the score. P-values were adjusted by the Benjamini-Hochberg (BH) method ^31^ for multiple hypotheses. For comparison with other driver gene prediction methods, we considered a gene to be significant (q≤0.1) if any of the OG, TSG, or driver scores were significant.

### Unexplained variability affects power and false positives

To evaluate effects of unexplained variability in the mutation rate on false positives and statistical power, we established two statistical models. The first model assumes a correctly estimated background mutation rate *μ* for a particular gene (binomial model) and the second model assumes that gene background mutation rate varies around *μ* (beta-binomial model). We used a binomial model previously developed for driver gene power analysis ^14^. The gene-specific mutation rate factor *F_g_* calculated by MutsigCV ^14^ was set to represent a gene at the 90^th^ percentile, given an exome-wide background mutation rate of *π* so that *μ* = *F_g_π* (*F_g_*= 3.9). Average gene length (L) was set to 1,500 bases and % mutations were assumed to be non-silent. Effective gene length was adjusted as *L_eff_* = ^3/4^ *L*. Gene background mutation rate was calculated using the total number of potentially mutated bases that could yield a non-silent mutation (*N_eff_*), which is the effective gene length multiplied by number of samples (S). A predicted driver was defined as a gene with significantly higher non-silent mutation rate per-base than that gene’s background mutation rate, where non-silent mutation rate per-base is and *r* is the fraction of samples with non-silent mutations in the gene above background. Exome-wide background mutation rates of (π=.5e-6, 3e-6, or 10e-6) were considered.

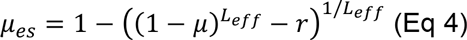

The beta-binomial was designed to model several levels of unexplained variability around *μ*. To parameterize the beta-binomial with low, medium and high variability levels, we used different coefficients of variation (CV) for the mutation rate (0.05, 0.1, 0.2). Beta-binomial *α* and *β* parameters were computed as

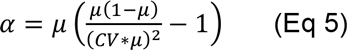

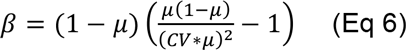

To compute the number of false positives expected from a binomial model when unexplained variability is present, we examined the probability that the number of mutations in a gene from a beta-binomial model (*K_beta-binomia_*) would meet or exceed the critical value (for a genome-wide significant driver gene at alpha = 5e-6) by the binomial *k’ _binomial_*.

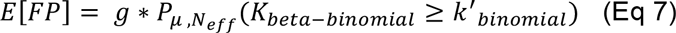

where g is the total number of human genes (assumed 18,500) and both models use the same mean mutation rate *μ* and total number of potentially mutated bases *N_eff_*.

We also modeled the effect of various levels of unexplained variability in mutation rate on the power to detect driver genes. We reproduced the binomial model power analysis of ^14^ to estimate the number of samples required for 90% power to detect genes in the 90th percentile of gene-specific background rate, with 2% mutation rate above background (r=0.02). Using Eq 5–6 to parameterize the beta-binomial model, we calculated the number of samples required for 90% power at a Bonferroni genome-wide significance level of 5e-6. Samples were iteratively added until there was greater than or equal to 90% probability that a driver gene with mutation rate *μ_es_* would be found significant. Using the jagged power curve for discrete data ^32^, we found the minimum number of samples required to achieve 90% power.

Next we evaluated the effects of unexplained variability on a ratiometric feature. As a proof of principle, we defined a simple ratiometric feature as the fraction of mutations in a gene having a specific non-silent mutation consequence type. As in the evaluation of mutation rate, we established two statistical models, assuming independence of mutation events. For the binomial model, the fraction is correctly estimated, and for the beta-binomial model the fraction varies around that estimate. A predicted driver gene was defined as having a significantly increased fraction with respect to the gene’s background. The fraction in a driver gene is then

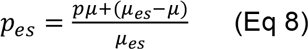

where *p* is the gene’s background fraction of the specific mutation consequence type and *μ* and *μ_es_* are calculated as in our above described analysis of mutation rate. In Figure 3B and 3C, *p* was set to 0.107 which is the fraction of inactivating mutations in our pan-cancer dataset.

As in the mutation rate analysis, we parameterized the beta-binomial with low, medium and high variability levels, using different coefficients of variation (CV) for the mutation rate (0.05, 0.1, 0.2). Beta-binomial *α* and *β* parameters were computed as in Eq 5 and Eq 6, but with μ replaced by *p*. Expected false positives were computed as in Eq 7, but with *μ* replaced by *p* and *N_eff_* -- the total number of potentially non-silent mutated bases -- replaced by the total number of expected non-silent or silent mutated bases

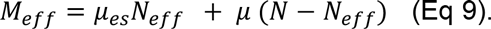

To estimate the effect of unexplained variability in the ratiometric feature on power to detect driver genes, we applied the same protocol used to assess unexplained variability in mutation rate on power. We estimated the minimum number of samples required for the same power and effect size used in the mutation rate analysis.

## Acknowledgments

Research funded by NCI F31CA200266 to CT, NCI 5U01CA180956-03 to RK and the Ludwig Foundation.

